# Karyorelict ciliates use an ambiguous genetic code with context-dependent stop/sense codons

**DOI:** 10.1101/2022.04.12.488043

**Authors:** Brandon Kwee Boon Seah, Aditi Singh, Estienne Carl Swart

## Abstract

In ambiguous stop/sense genetic codes, the stop codon(s) not only terminate translation but can also encode amino acids. Such codes have evolved at least four times in eukaryotes, twice among ciliates (*Condylostoma magnum* and *Parduczia* sp.). These have appeared to be isolated cases whose next closest relatives use conventional stop codons. However, little genomic data have been published for the Karyorelictea, the ciliate class that contains *Parduczia* sp., and previous studies may have overlooked ambiguous codes because of their apparent rarity. We therefore analyzed single-cell transcriptomes from four of the six karyorelict families to determine their genetic codes. Reassignment of canonical stops to sense codons was inferred from codon frequencies in conserved protein domains, while the actual stop codon was predicted from full-length transcripts with intact 3’-untranslated regions (3’-UTRs). We found that all available karyorelicts use the *Parduczia* code, where canonical stops UAA and UAG are reassigned to glutamine, and UGA encodes either tryptophan or stop. Furthermore, a small minority of transcripts may use an ambiguous stop-UAA instead of stop-UGA. Given the ubiquity of karyorelicts in marine coastal sediments, ambiguous genetic codes are not mere marginal curiosities but a defining feature of a globally distributed and diverse group of eukaryotes.

## Introduction

In addition to the “standard” genetic code used by most organisms, there are numerous variant codes across the tree of life, and new ones continue to be discovered [1–3]. The differences between codes lie in which amino acids are coded by which codon, as well as which codons are used to start and terminate translation (stop codons). Much of the variation is concentrated in a small number of codons, particularly the canonical stop codons UAA, UAG, and UGA, which have repeatedly been reassigned to encode amino acids. The most striking variants are ambiguous codes where one codon can have multiple meanings. The outcome during translation can be stochastic, such as in stop codon readthrough [4], or translation of CUG as either leucine or serine by *Candida* spp. [5]. Alternatively, they can be context-dependent, such as UGA encoding selenocysteine only in selenoproteins [6], meaning that the translation system is able to interpret the codon correctly as either an amino acid or a stop.

Other context-dependent stop/sense codes have been discovered where all the stop codons used by the cell are potentially also sense codons. These have evolved independently several times among the eukaryotes [7–10]: parasitic trypanosomes of the genus *Blastocrithidia* (three different species) use UAA and UAG to encode stop/glutamate (NCBI Genetic Codes ftp.ncbi.nih.gov/entrez/misc/data/gc.prt, table 31); a strain of the marine parasitic dinoflagellate *Amoebophrya* and a marine karyorelict ciliate, *Parduczia* sp., have convergently evolved to use UGA for stop/tryptophan (table 27); and the marine heterotrich ciliate *Condylostoma magnum* uses UGA for stop/tryptophan and UAA/UAG for stop/glutamine (table 28).

The ciliates are a clade with an unusual propensity for variant genetic codes [11]. At least eight different nuclear genetic codes are used by ciliates [10], including some of the first examples of variant codes documented in nuclear genomes [12–16]. At first glance, organisms that use these ambiguous stop/sense codes appear to be isolated single species or strains embedded among relatives with conventional codes. For example, other heterotrichs related to *Condylostoma* use the standard code (e.g. *Stentor*) or the *Blepharisma* code. Additionally, a previous survey of genetic codes across the ciliate tree, including numerous uncultivated heterotrichs and karyorelicts, did not report any new examples of organisms that use ambiguous stop/sense codes, nor appeared to have accounted for such a possibility in their methods [17]. However our own preliminary studies appeared to contradict this, finding other karyorelicts that use the same genetic code as *Parduczia*.

The karyorelicts are a class-level taxon within the ciliates, and sister group to the heterotrichs. Unlike other ciliates, the somatic nuclei (macronuclei) of karyorelicts do not divide but must differentiate anew from germline nuclei (micronuclei) every time, even during vegetative division [18]. They are globally distributed and commonly encountered in the sediment interstitial habitat of marine coastal environments [19]. At least ∼150 species have been formally described but this is believed to be a severe underestimate of the true diversity [20,21], and they are also poorly represented in sequence databases.

We therefore sequenced additional karyorelict transcriptomes and reanalyzed published data to assess whether karyorelicts other than *Parduczia* could be using ambiguous genetic codes.

## Results

Ten new single-cell RNA-seq libraries from karyorelicts and heterotrichs were sequenced in this study, representing interstitial species from marine sediment at Roscoff, France. These were analyzed alongside 33 previously published RNA-seq libraries (doi:10.17617/3.XWMBKT, Table S1). After filtering for quality and sufficient data, 25 transcriptome assemblies (of which 15 were previously published) were used to evaluate stop codon reassignment, vs. 26 assemblies (16 previously published) for inferring the actual stop codon(s) (Appendix).

### Reassignment of all three canonical stop codons to sense codons in karyorelicts

Codon frequencies in protein-coding sequences were calculated from sequence regions that aligned to conserved Pfam domains, in transcripts with poly-A tails. Transcriptomes and genomic coding sequences (CDSs) from ciliates with known genetic codes were used as a comparison to estimate the false positive rate of stop codons being found in these alignments, e.g. because of misalignments, misassembly, or pseudogenes.

Among karyorelicts, all three canonical stop codons (UAA, UAG, UGA) were observed in conserved protein domains, with frequencies between 0.08-2.9%, which fell within the range of codon frequencies observed for unambiguous sense codons in other ciliates where the genetic code is known (0.03-6.8%, excluding the outlier CGG in *Tetrahymena thermophila* with only 0.003%). This range was also similar to frequencies of the ambiguous stops in *Parduczia* and the heterotrich *Condylostoma* (Figure 1A). UGA was generally less frequent than UAA/UAG in all karyorelicts, but the frequencies varied between taxa, reflecting their individual codon usage biases or which genes are assembled in the transcriptome because of sequencing depth. UGA was the least-frequent codon in most Trachelocercidae and Geleiidae, but was more frequent in Loxodidae and Kentrophoridae than some other codons, especially C/G-rich ones like CGG (Figure 1A). Nonetheless, frequencies of the UGA codon in karyorelicts were all still one to two orders of magnitude higher than the observed frequencies of in-frame actual stops from other ciliate species in the reference set.

**Figure 1.**
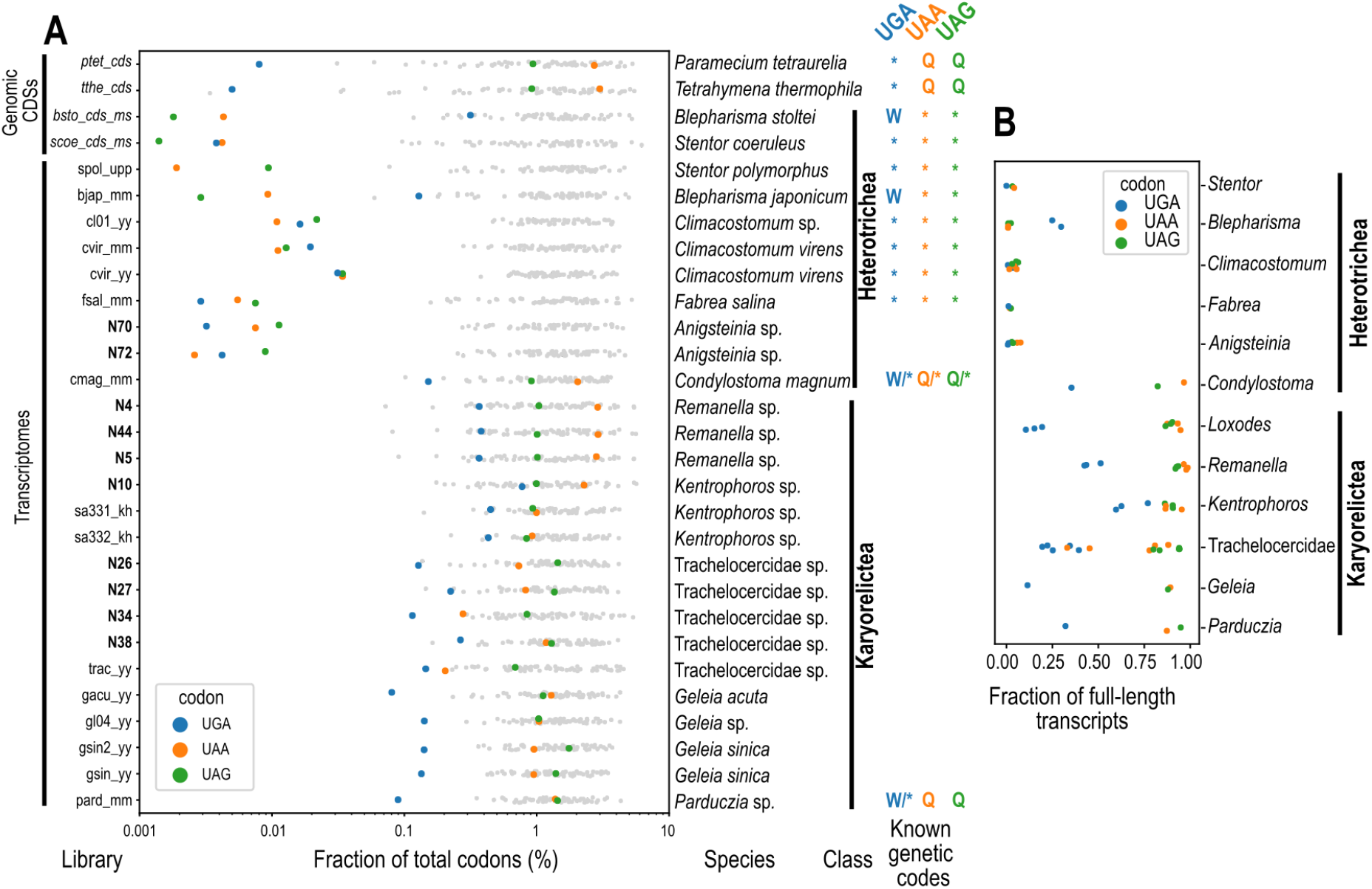
(**A**) Codon frequencies of canonical stop codons (UGA: blue, UAA: orange, UAG: green) and other codons (gray) in conserved protein domains found by hmmscan search in six-frame translations of transcriptome assemblies (doi:10.17617/3.XWMBKT, Table S1) or genomic CDSs (doi:10.17617/3.XWMBKT, Table S2) vs. Pfam. Names of libraries from this study are highlighted in bold. Assignments of canonical stops for organisms with known genetic codes follow ref. [10]. (**B**) Fraction of full-length transcripts that have at least one canonical stop codon in the putative coding region, grouped by genus (except Trachelocercidae, where classification was unclear).

In-frame UGAs were found in 10.5 to 76.9% of transcripts with putative coding regions predicted by full-length Blastx hits per karyorelict library (Figure 1B). This frequency verified that in-frame UGAs were not concentrated in a small fraction of potentially spurious sequences but in fact found in many genes. Conserved “marker” genes that were generally expected to be present in ciliate genomes (BUSCO orthologs, Alveolata marker set, [22]) also contained in-frame UGAs. The karyorelict transcriptome assemblies were relatively incomplete, with 1.8% to 20.5% (median 12.0%) estimated completeness based on the BUSCO markers, and a total of 91 of 171 BUSCO orthologs were found in these assemblies (Figure 2A). Nonetheless, 46 BUSCO orthologs from 14 karyorelict assemblies were found with in-frame UGAs in conserved alignment positions (e.g. Figure 2B, 2C), verifying that they are not limited to poorly characterized or hypothetical proteins.

**Figure 2.**
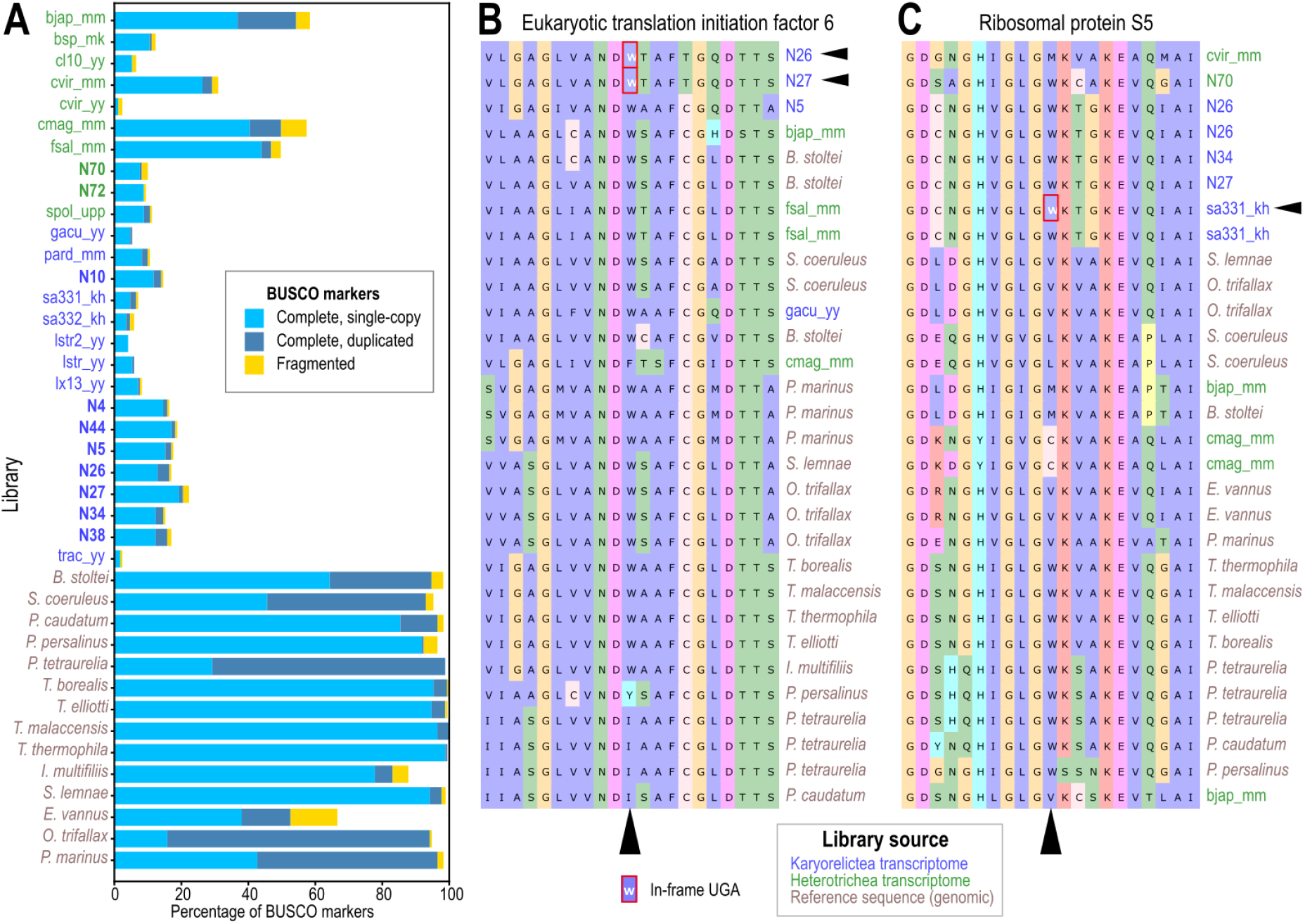
In-frame coding UGAs in conserved marker genes. (**A**) Completeness estimates of heterotrich and karyorelict transcriptomes (library names in green and blue respectively), compared with genomic reference sequences from other ciliates (doi:10.17617/3.XWMBKT, Table S3); BUSCO Alveolata marker set. (**B, C**) Two examples of alignments (excerpts) for conserved orthologous protein-coding genes (orthologs 20320at33630 and 23778at33630), which contain in-frame UGAs translated as W in karyorelict sequences, flanked by conserved alignment blocks.

In comparison, the heterotrich *Anigsteinia*, for which two new sequence libraries were also produced and which was found in the same habitats as karyorelicts, had in-frame frequencies of ≤0.011% for all three canonical stop codons, which were comparable to frequencies of the known stop codons in *Blepharisma* (UAA, UAG) and *Stentor* (UAA, UAG, UGA) (max. 0.09%). Hence *Anigsteinia* probably does not have ambiguous sense/stop codons.

All karyorelicts had the same inferred amino acid reassignments for the three canonical stops: glutamine (Q) for UAA and UAG, and tryptophan (W) for UGA (Figure 3), matching previous predictions for *Parduczia* sp. and *Condylostoma magnum* [9,10].

**Figure 3.**
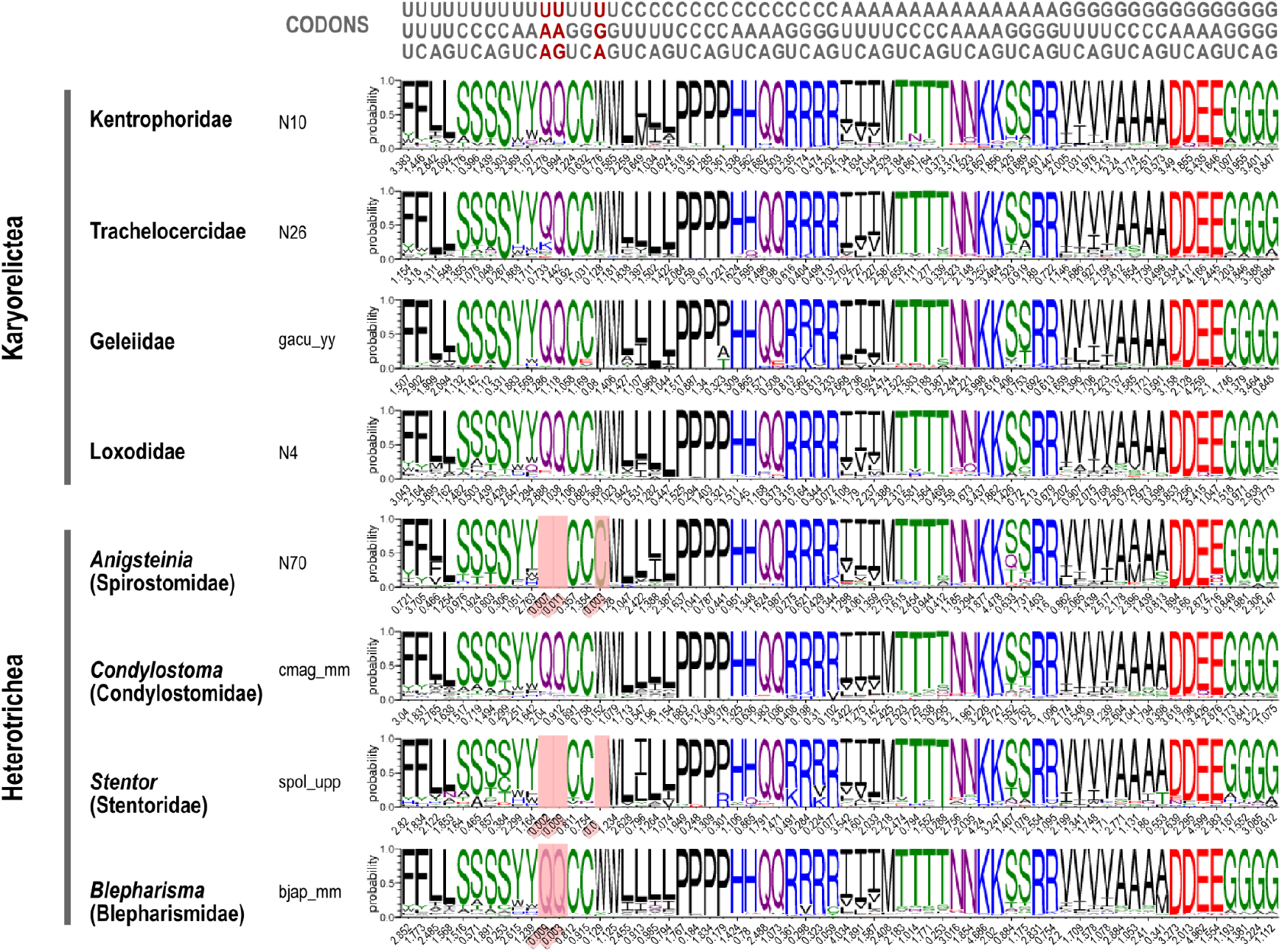
Weblogos representing the likely amino acid assignment of each codon in selected libraries (library with most coverage per taxon of interest). Heights of each letter represent the relative frequencies (all scaled to 100%) of each amino acid in conserved residues aligning to that codon. The observed codon frequency (in %) is indicated below. Codons with frequencies <0.02% are highlighted in red, representing either non-ambiguous stops or unassigned codons. Assignment of cysteine (C) for UGA in *Anigsteinia* is based on only 16 alignments, of which 14 are to a likely selenoprotein (Pfam domain GSHPx); assignment of glutamine (Q) for UAA and UAG in *Blepharisma* may represent recent paralogs or translational readthrough.

### Stop codons in karyorelicts and heterotrichs

Frequency of a codon in coding regions can be used to infer if it is a sense codon but not whether it can terminate translation, especially for ambiguous codes where codons that can terminate translation also frequently appear in coding sequences. Therefore we used full length transcripts with both a high quality Blastx alignment to a reference protein and a poly-A tail to predict the likely stop codon(s) used in each sample. To avoid double counting, only one isoform was used per gene. We assumed that the true stop codon(s) were one or more of the three canonical stops UGA, UAA, UAG, and that if a contig has a high quality Blastx hit to a reference protein sequence, the true stop should lie somewhere between the last codon at the 3’ end of the hit region and the beginning of the poly-A. We reasoned that if the true stop codon set was used for annotation, (i) the number of transcripts without a putative true stop should be minimized; (ii) the variance of the 3’-untranslated region (3’-UTR) length should also be minimized because ciliate 3’-UTRs are known to be short (mostly <100 bp); and (iii) if there was more than one stop codon, the length distributions of the putative 3’-UTRs for each stop codon should be centered on the same value.

With these criteria, the candidate stop codons for karyorelicts could be narrowed to two possibilities: UGA alone or UGA + UAA. If only UGA was permitted as a stop codon, 84-98% of transcripts per library had a putative true stop, but if both UGA and UAA were permitted as stop codons, the proportion was over 98% (Figure 4A). Permitting both UGA+UAA as stops in karyorelicts resulted in a higher variance in 3’-UTR lengths compared to permitting only UGA. Although this was contrary to criterion (ii) above, we judged that this metric was not as useful in deciding whether UAA was also a stop codon, because the difference was small, and transcripts with putative UAA stops were relatively few (Figures 4B, 4C). Both karyorelicts and heterotrichs in this study had short and narrowly distributed 3’-UTR lengths (median 28 nt, interquartile range 18 nt) (Figure 4C). The heterotrichs were shortest overall, with median lengths per taxon between 21 nt (*Condylostoma*) and 26 nt (*Stentor*), followed by the karyorelict families Trachelocercidae (33 nt), Geleiidae (31 nt), Kentrophoridae (37 nt), and Loxodidae (43 nt).

**Figure 4.**
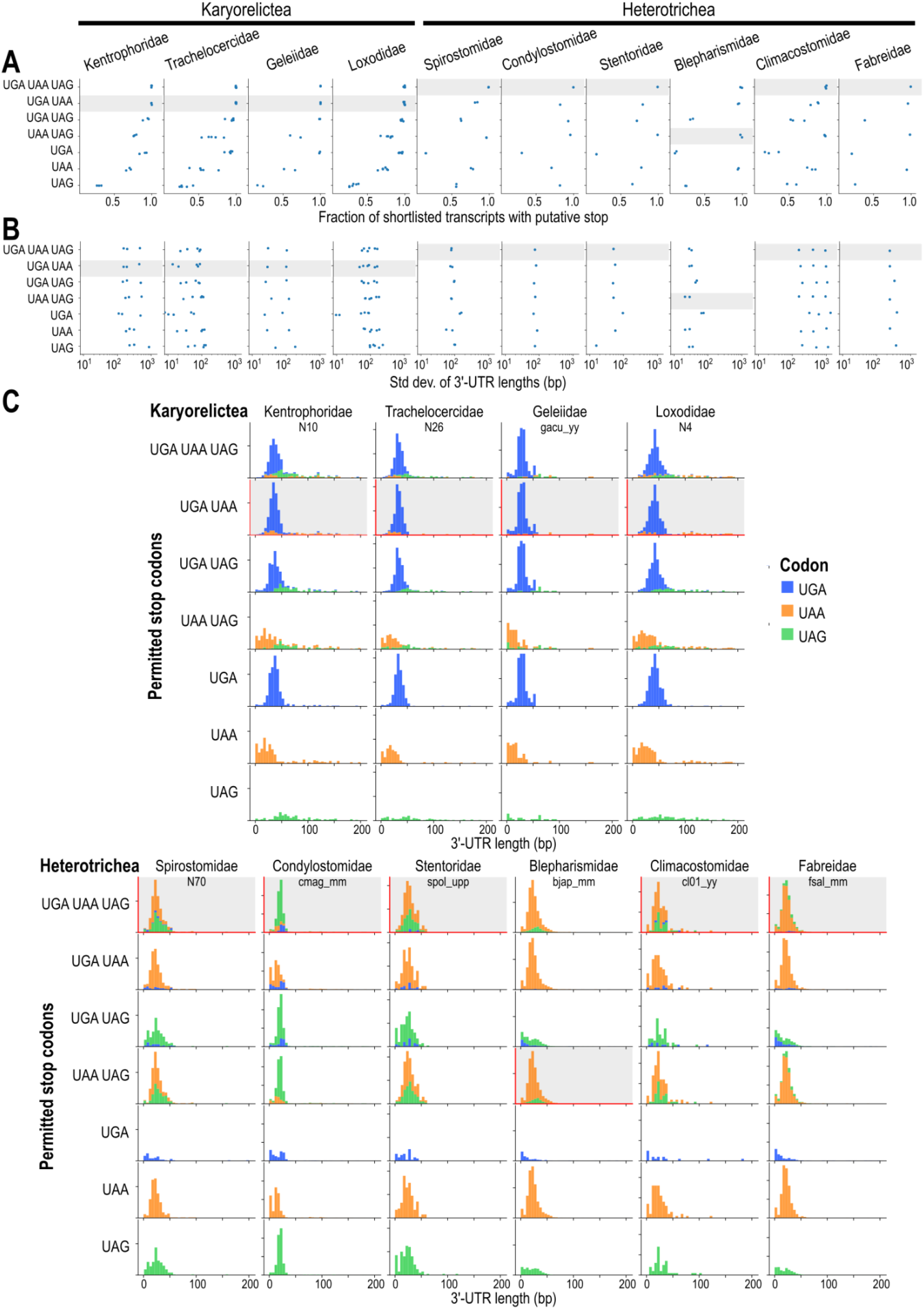
Effect of different stop codon combinations on assembly metrics. Predicted stop codon usage for each taxon from this study or previous publications highlighted in gray. (**A**) Strip plots for the fraction of full length contigs per transcriptome that have a putative stop codon from that specific combination (rows), i.e. in-frame, downstream of full-length Blastx hit vs. reference, and upstream of poly-A tail. Each point corresponds to one transcriptome assembly, grouped by taxonomic family (columns). (**B**) Scatterplots for standard deviation of 3’-UTR lengths. (**C)** Histograms for 3’-UTR lengths, colored by putative stop codon (UGA: blue, UAA: orange, UAG: green), one representative library per family.

In previous analyses of the ambiguous stop codons in *Condylostoma* and *Parduczia*, a distinct depletion of in-frame coding “stop” codons immediately upstream of the actual terminal stop was observed [10]. We could reproduce this depletion of all three canonical stops in *Condylostoma* and of UGA in *Parduczia*, about 10 to 20 codon positions before the putative terminal stop, in our reanalysis of the same data (Figure 5A). For the karyorelicts, if only UGA was permitted as a stop codon, we observed depletion of coding-UGA but also of coding-UAAs before the terminal stop-UGA (Figure 5B). If UGA + UAA were permitted as stops, the depletion of coding-UGA before terminal stops was still observed, and the depletion of coding-UAA was even more pronounced (Figure 5C). Unfortunately, there were only a limited number of full-length karyorelict transcripts with putative stop-UAAs (max. 47 contigs per library). We therefore concluded that UGA is the predominant stop codon in karyorelicts, but UAA may also function as a stop codon for about 1-10% of transcripts.

**Figure 5.**
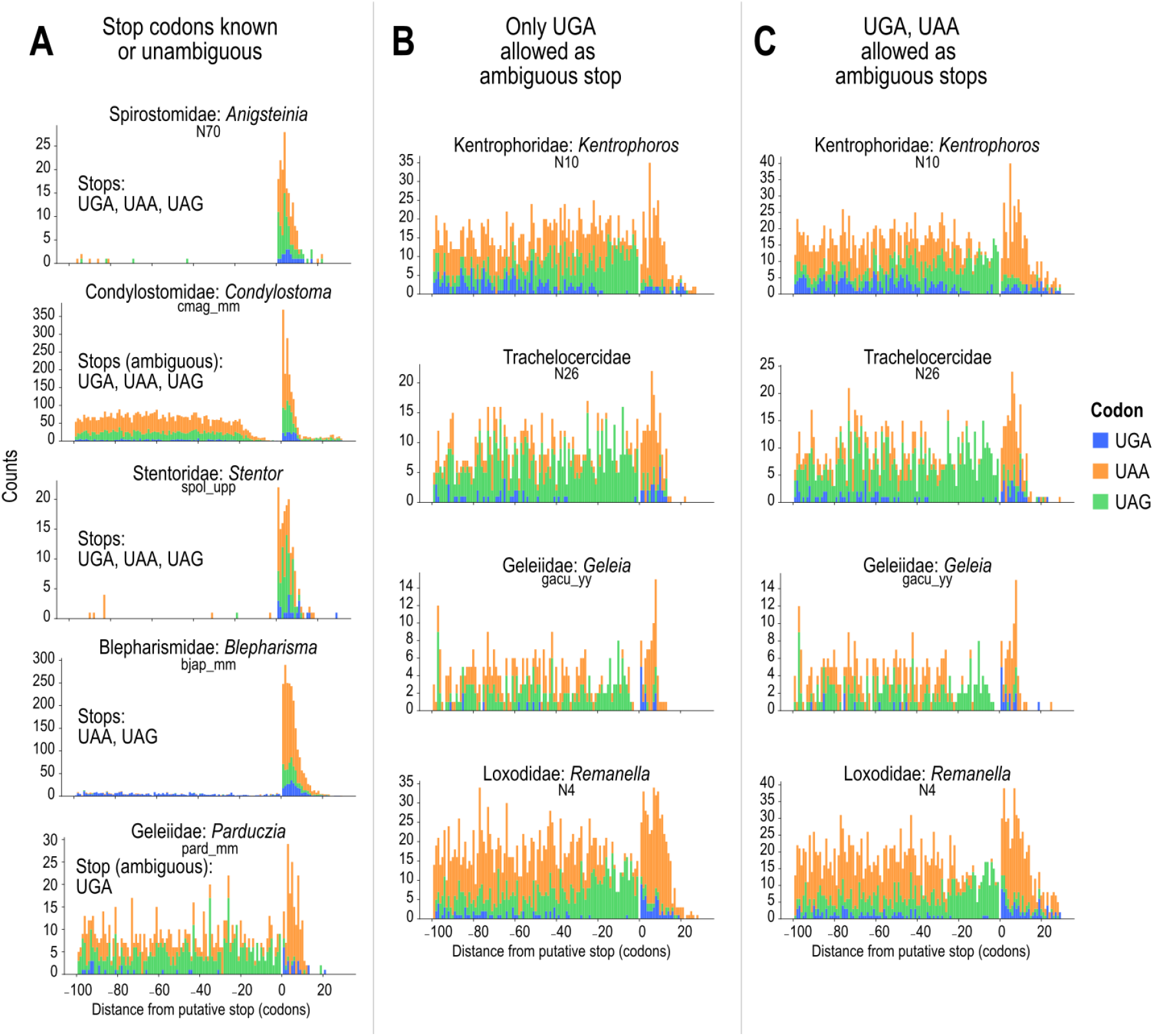
Depletion of in-frame coding “stop” codons in the coding sequence (negative coordinates) immediately before the putative true stop codon (position 0) and their enrichment in the 3’-UTR (positive coordinates). Representative library with highest number of assembled full length contigs chosen per taxon. (**A**) Codon counts for UGA (blue), UAA (orange), and UAG (green) before and after putative true stop in *Condylostoma magnum* (uses all three as ambiguous stops), and three heterotrichs with unambiguous stops. (**B**) Codon counts for karyorelicts if only UGA is permitted as a stop codon. (**C**) Codon counts for karyorelicts if both UGA and UAA are permitted as stop codons.

UAA and UAG were predicted as stop codons of *Anigsteinia* (Spirostomidae), consistent with their near-absence from coding regions in this genus (see above, Figure 1A). UGA was not only near-absent from coding regions, but also rarely encountered as a putative stop codon, although it was not uncommon in 3’-UTRs. Similar rarity of UGAs as putative stops was also observed in *Stentor* and other heterotrichs that are said to use the standard code. Either (i) these heterotrichs use the standard genetic code with all three canonical stop codons but a strong bias against using UGA for stop, or (ii) UGA is an unassigned codon in these organisms.

## Discussion

We have found evidence that the codon UGA is used as both a stop codon and to code for tryptophan by karyorelictean ciliates. The taxa sampled represent four of the six families of karyorelicts: Loxodidae, Trachelocercidae, Geleiidae, and Kentrophoriidae. When this distribution of genetic codes is mapped to an up-to-date phylogeny [20], we can infer that the ambiguous code formerly reported only for *Parduczia* sp. (Geleiidae) among ciliates was actually acquired at the root of the karyorelict clade (Figure 6).

**Figure 6.**
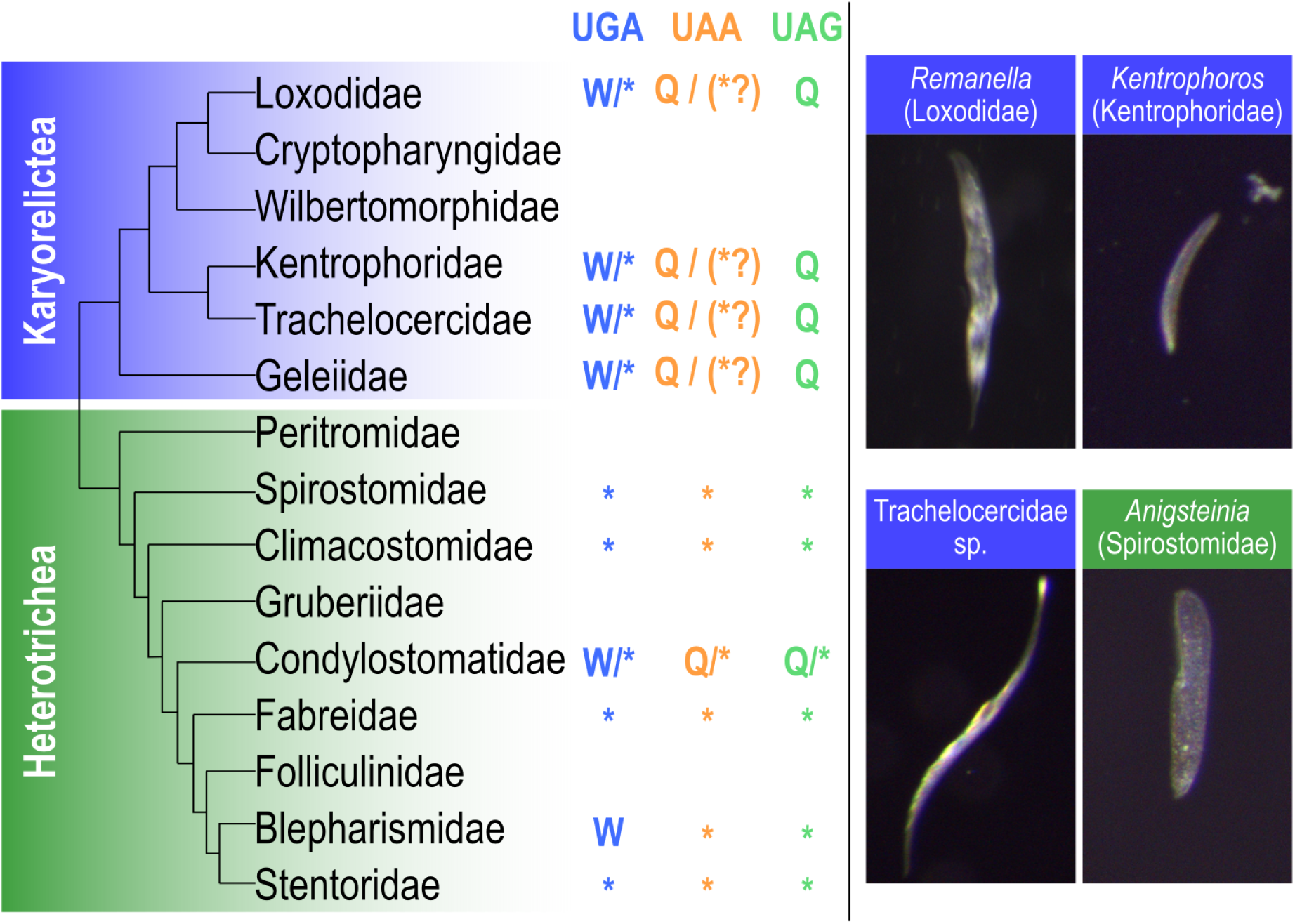
Genetic code diversity among karyorelict and heterotrich ciliates. (**Left**) Diagrammatic karyorelict + heterotrich tree with predicted stop codon reassignments mapped to each family. Subtree topologies are from refs. [20] and [24] respectively. Branch lengths are not representative of evolutionary distances. (**Right**) Photomicrographs of ciliates (incident light) collected in this study from Roscoff, France; height of each panel 50 µm.

Available data for *Cryptopharynx* (Karyorelictea: Cryptopharyngidae) were not conclusive. The canonical stop codons had frequencies between 0.02 and 0.07%, lower than for other karyorelicts, but higher than true stop codons, but Cryptopharyngidae was represented by a single library that had high contamination from other eukaryotes (Appendix) and there were too few high-confidence, full length transcripts for a reliable conclusion on its genetic code. No sequence data beyond rRNA genes were publicly available for the remaining family, the monotypic Wilbertomorphidae, whose phylogenetic position in relation to the other karyorelicts is unclear because of long branch lengths, and which has to our knowledge only been reported once [23].

Ambiguous stop/sense codes are hence not just isolated phenomena, but are used by a major taxon that is diverse, globally distributed, and common in its respective habitats. In contrast, the heterotrichs, which constitute the sister group to Karyorelictea and are hence of the same evolutionary age, use at least three different genetic codes, including one with ambiguous stops (Figure 6). If organisms with ambiguous codes were isolated single species whose nearest relatives have conventional stops, as appears to be the case for *Blastocrithidia* spp. and *Amoebophrya* sp., we might conclude that these are uncommon occurrences that do not persist over longer evolutionary time scales. However, the karyorelict crown group diversified during the Proterozoic (posterior mean 455 Mya) and the stem split from the Heterotrichea even earlier, in the Neo-Proterozoic [24].

This study has benefited from several technical improvements. A highly complete, contiguous genome assembly with gene predictions is now available for the heterotrich *Blepharisma stoltei* [25]. Because *Blepharisma* is more closely related to the karyorelicts than other ciliate model species, which are mostly oligohymenophorans and spirotrichs, it improved the reference-based annotation of the assembled transcriptomes. Single-cell RNA-seq libraries in this study were also sequenced to a greater depth, with a lower fraction of contamination from rRNA, and hence yielded more full length mRNA transcripts for analysis.

One proposed mechanism for how the cell correctly recognizes whether an ambiguous codon is coding or terminal is based on the proximity of translation stops to the poly-A tail of transcripts. In this model, tRNAs typically bind more efficiently to in-frame coding “stops” than eukaryotic translation release factor 1 (eRF1), hence allowing these codons to be translated. At the true termination stop codon, however, the binding of eRF1 can be stabilized by interactions with poly-A interacting proteins like PABP bound to the nearby poly-A tail, allowing it to outcompete tRNAs and hydrolyze the peptidyl-tRNA bond [10,26]. Consistent with this model, we found that karyorelict 3’-UTRs are also relatively short, and that in-frame UGAs are depleted immediately before the putative true stop codon. Nonetheless, karyorelict 3’-UTRs are actually about 10 nt longer on average than those of heterotrichs.

Our results raised the possibility that UAA is also used as an ambiguous stop codon for ∼1-10% of karyorelict transcripts, in addition to the main stop codon UGA. eRF1 may retain a weak affinity for UAA, and recognize UAA for terminating translation albeit with lower efficiency. In *Blepharisma japonicum*, where UAA and UAG are non-ambiguous stops and UGA encodes tryptophan (albeit at low frequency, 0.13%), heterologously expressed eRF1 could still recognize all three codons in an in vitro assay, although efficiency of peptidyl-tRNA hydrolysis was lower with UGA than for UAA and UAG [27]. In species with non-ambiguous stop codon reassignment, the effect of such “weak” ambiguity on the total pool of translated protein may be negligible, but it shows that there is a latent potential that could account for the repeated evolution of stop codon reassignments in ciliates. Furthermore, UAAs were even more abundant than UGAs in ciliate 3’-UTRs, which can be attributed to the low GC% of 3’-UTRs compared to coding sequences; other A/U-only codons were also enriched in 3’-UTRs. Therefore, UAAs in the 3’-UTRs of karyorelicts may be a “backstop” mechanism that prevents occasional stop-codon readthrough, as proposed for tandem stop codons (TSCs) in other species with reassigned stop codons [28]. In the minority of transcripts where in-frame stop-UGA is absent, the backstop may be adequate to terminate translation before the poly-A tail and produce a functional protein most of the time. To verify our predictions that UGA is the main stop codon and UAA a lower-frequency alternative stop, ribosome profiling and mass spectrometry detection of peptide fragments corresponding to the expected 3’-ends of coding sequences, e.g. as performed on *Condylostoma* [10], are the most applicable experimental methods. If a karyorelict species can be developed into a laboratory model amenable to genetic transformation, manipulation of the 3’-UTR length and sequence would allow us to test the “backstop” hypothesis directly and tease apart the factors contributing to translation termination in these organisms.

What selective pressures might favor the evolution and maintenance of an ambiguous genetic code? One possibility is that context-dependent sense/stop codons confer mutational robustness by eliminating substitutions that cause premature stop codons. Ambiguous codes do not appear to be linked to a specific habitat: *Blastocrithidia* spp. and *Amoebophrya* sp. are both parasites of eukaryotic hosts, but of insects and free-living dinoflagellates respectively; whereas the karyorelict ciliates and *Condylostoma* are both found in marine interstitial environments, but live alongside other ciliates that have conventional codes, such as *Anigsteinia*. Having short 3’-UTRs may predispose ciliates to adopt ambiguous codes by facilitating interactions between eRF1 and PABPs that could enable stop recognition, but other factors, including simply contingent evolution, appear to have led to their evolution because the 3’-UTRs of ciliates with conventional stop codons are also comparably short, particularly among the heterotrichs.

Any adaptationist hypothesis for alternative and ambiguous codes will have to contend with the existence of related organisms with conventional codes that have similar lifestyles. Furthermore, once a stop codon has been reassigned to sense, it becomes increasingly difficult to undo without the deleterious effects of premature translation termination, and may function like a ratchet. Like the origins of the genetic code itself [29], we may have to be content with the null hypothesis that they are “frozen accidents” that reached fixation stochastically, and which are maintained because they do not pose a significant selective disadvantage.

## Materials and Methods

### Sample collection

Surface sediment was sampled in September 2021 from two sites in the bay at Roscoff, France when exposed at low tide. Site A: shallow swimming enclosure, 48.72451 N, 3.992294 W; Site B: adjacent to green algae tufts near freshwater outflow, 48.716169 N, 3.995626 W. Upper 1-2 cm of sediment was skimmed into glass beakers, and stored under local seawater until use. Interstitial ciliates were collected by decantation: a spoonful of sediment was stirred in seawater in a beaker. Sediment particles were briefly allowed to settle out, and the overlying suspended organic material was decanted into Petri dishes. Ciliate cells were preliminarily identified by morphology under a dissection microscope and picked by pipetting with sterile, filtered pipette tips. Selected cells were imaged with incident light under a stereo microscope (Olympus SZX10, Lumenera Infinity 3 camera).

NEBNext cell lysis buffer (NEB, E5530S) was premixed and filled into PCR tubes; per tube: 0.8 µL 10x cell lysis buffer, 0.4 µL murine RNAse inhibitor, 5.3 µL nuclease-free water. Picked ciliate cells were transferred twice through filtered local seawater (0.22 µm, Millipore SLGP033RS) to wash, then transferred with 1.5 µL carryover volume to 6.5 µL of cell lysis buffer (final volume 8 µL), and snap frozen in liquid nitrogen. Samples were stored at -80 °C before use.

### Single-cell RNAseq sequencing

Samples collected in cell lysis buffer (doi:10.17617/3.XWMBKT, Table S1) were used for RNAseq library preparation with the NEBNext Single Cell / Low Input RNA Library Prep Kit for Illumina (NEB, E6420S), following the manufacturer’s protocol for single cells, with the following parameters adjusted: 17 cycles for cDNA amplification PCR, cDNA input for library enrichment normalized to 3 ng (or all available cDNA used for libraries where total cDNA was <3 ng), 8 cycles for library enrichment PCR. Libraries were dual-indexed (NEBNext Dual Index Primers Set 1, NEB E7600S), and sequenced on an Illumina NextSeq 2000 instrument with P3 300 cycle reagents, with target yield of 10 Gbp per library.

### RNA-seq library quality control and transcriptome assembly

Previously published karyorelict transcriptome data [17,30–32] were downloaded from the European Nucleotide Archive (ENA) (doi:10.17617/3.XWMBKT, Table S1). Contamination from non-target organisms was evaluated by mapping reads to an rRNA reference database and summarizing the hits by taxonomy. Although RNAseq library construction enriches mRNAs using poly-A tail selection, there is typically still sufficient rRNA present in the final library to evaluate the taxonomic composition of the sample. All RNAseq read libraries (newly sequenced and previously published) were processed with the same pipeline: The taxonomic composition of each library was evaluated by mapping 1 M read pairs per library against the SILVA SSU Ref NR 132 database [33], using phyloFlash v3.3b1 [34]. Newly sequenced libraries were assigned to a genus or family using the mapping-based taxonomic summary, or full-length 18S rRNA gene if it was successfully assembled.

Reads were trimmed with the program bbduk.sh (http://sourceforge.net/projects/bbmap/, BBmap v38.22) to remove known adapters (right end) and low-quality bases (both ends), with minimum Phred quality 24 and minimum read length 25 bp. Trimmed reads were then assembled with Trinity v2.12.0 [35] using default parameters. Assembled contigs were aligned against the *Blepharisma stoltei* ATCC 30299 proteome [25] with NCBI Blastx v2.12.0 [36] using the standard genetic code and E-value cutoff 10-20, parallelized with GNU Parallel [37].

Morphological identifications of the newly collected samples were verified with 18S rRNA sequences from the Trinity transcriptome assemblies (Appendix). rRNA sequences were annotated with barrnap v0.9. 18S rRNA sequences ≥80% of full length were extracted, except for two libraries (N4, N26) where the longest sequences were <80% and for which the two longest 18S rRNA sequences were extracted instead. For comparison, reference sequences for Karyorelictea and Heterotrichea above 1400 bp from the PR2 database v4.14.0 [38] were used. Representative reference sequences were chosen by clustering at 99% identity with the cluster_fast method using Vsearch v2.13.6 [39]. Extracted and reference sequences were aligned with MAFFT v7.505 [40]. A phylogeny (Figure S3) was inferred from the alignment with IQ-TREE v2.0.3 [41], using the TIM2+F+I+G4 model found as the best-fitting model by ModelFinder [42]. Alignment and tree files are available from doi:10.17617/3.QLWR38. 18S rRNA sequences were deposited in the European Nucleotide Archive under accessions OX095806-OX095846.

Read pre-processing, quality control, and assembly were managed with a Snakemake v6.8.1 [43] workflow (https://github.com/Swart-lab/karyocode-workflow, archived at doi:10.5281/zenodo.6647650). Scripts for data processing described below were written in Python v3.7.3 using Biopython v1.74 [44], pandas v0.25.0 [45], seaborn v0.11.0 [46] and Matplotlib v3.1.1 [47] libraries unless otherwise stated.

### Prediction of stop codon reassignment to sense

Only contigs with poly-A tails ≥7 bp were used for genetic code prediction, to exclude potential bacterial contaminants, especially because several species (*Kentrophoros* spp., *Parduczia* sp., Appendix) are known to have abundant bacterial symbionts. Presence and lengths of poly-A tails in assembled transcripts were evaluated with a Python regular expression. Library preparation was not strand-specific, hence contigs starting with poly-T were reverse-complemented, and contigs with both a poly-A tail and a poly-T head (presumably fused contig) were excluded.

Codon frequencies and their corresponding amino acids were predicted with an updated version of PORC (v2.1, https://github.com/Swart-lab/PORC, archived at doi:10.5281/zenodo.6784075; managed with a Snakemake workflow, https://github.com/Swart-lab/karyocode-analysis-porc, archived at doi:10.5281/zenodo.6647652); the method has been previously described [10,48]. Briefly: a six-frame translation was produced for each contig in the transcriptome assembly, and searched against conserved domains in the Pfam-A database v32 [49] with hmmscan from HMMer v3.3.2 (http://hmmer.org/). Overall codon frequencies were counted from alignments with E-value ≤ 10-20. To ensure that there was sufficient data underlying the codon and amino acid frequencies, only those libraries with at least 100 observations for each of the coding codons in the standard genetic code were used for comparison of codon frequencies and for prediction of amino acid assignments.

Frequencies of amino acids aligning to a given codon were counted from columns where the HMM model consensus was ≥50% identity in the alignment used to build the model (upper-case positions in the HMM consensus). Sequence logos of amino acid frequencies per codon for each library were drawn with Weblogo v3.7.5 [50].

In addition to the transcriptomes, genomic CDSs of selected model species with different genetic codes [25,51–55] were also analyzed with PORC to obtain a reference baseline of coding-codon frequencies (doi:10.17617/3.XWMBKT, Table S2). These model species have non-ambiguous codes so they were not expected to have stop codons in the CDSs, except for the terminal stop.

### Prediction of coding frame in full-length transcripts

“Full-length” transcripts (with poly-A tail, intact 3’-UTR, and complete coding sequence) were desirable to predict the stop codon, characterize 3’-UTR metrics, and verify genetic code predictions. Contigs were therefore filtered with the following criteria: (i) poly-A tail ≥7 bp, criterion following [10], (ii) contig contains a Blastx hit vs. *Blepharisma stoltei* protein sequence with E-value ≤10^−20^ and where the alignment covers ≥80% of the reference *B. stoltei* sequence, (iii) both poly-A tail and Blastx hit agree on the contig orientation. For contigs with multiple isoforms assembled by Trinity, the isoform with the longest Blastx hit was chosen; in case of a Blastx hit length tie, then the longer isoform was chosen. Only libraries with >100 assembled “full-length” transcripts were used for downstream analyses (Appendix).

### Metrics for evaluating potential stop codon combinations

For each of the 7 possible combinations of the 3 canonical stop codons (UGA, UAA, UAG), we treated the first in-frame stop downstream of the Blastx hit in each full-length transcript (including the last codon of the hit) as the putative stop codon, and recorded the number of full-length transcripts with a putative stop, the length of the 3’-UTR (distance from stop to beginning of the poly-A tail), as well as the codon frequencies for each position from 150 codons upstream of the putative stop to the last in-frame three-nucleotide triplet before the poly-A tail.

### Delimitation of putative coding sequences using Blastx hits

The start codon was more difficult to evaluate because the 5’ end of the transcript may not have been fully assembled, and there was no straightforward way to recognize its boundaries, unlike the 3’-poly-A tail. We used the following heuristic criteria to define the start of the CDS: first in-frame ATG upstream of the Blastx hit (including first codon of the hit), or first in-frame stop codon encountered upstream (to avoid potential problems with ORFs containing in-frame stops), whichever comes first. Otherwise, the transcript was assumed to be incomplete at the 5’-end and simply truncated with the required 1 or 2 bp offset to keep the CDS in frame.

### Verification of in-frame UGAs in conserved marker genes

Full-length CDSs (see above) were translated with the karyorelict code (NCBI table 27). Conserved marker genes were identified with BUSCO v5.2.2 (protein mode, alveolata_odb10 marker set) [22], managed with a Snakemake workflow (https://github.com/Swart-lab/karyocode-analysis-busco, archived at doi:10.5281/zenodo.6647679). Markers for additional ciliate species where relatively complete genome assemblies and gene predictions were available were also identified (doi:10.17617/3.XWMBKT, Table S3) [52,54,56–63]. For each BUSCO marker, the ciliate homologs were aligned with Muscle v3.8.1551 [64]. Alignment columns corresponding to in-frame putatively coding UGAs of karyorelict sequences were identified. These positions were considered to be conserved if ≥50% of residues were W or another aromatic amino acid (Y, F, or H).

## Acknowledgements

Version 4 of this preprint has been peer-reviewed and recommended by Peer Community In Genomics (https://doi.org/10.24072/pci.genomics.100019). We thank S. Booker, C. Cabresin, E. Bourrigaud, and R. Garnier for facilitating our research visit to the Station Biologique Roscoff; H. Budde for sequencing; H. Gruber-Vodicka for loan of equipment; A. Peters for helpful suggestions and cake; A. Lemper for administrative assistance; Y. Shulgina for thoughtful feedback on the manuscript; the recommender and peer reviewers for PCI Genomics for their reviews; and members of the Swart Lab for helpful feedback and support.

## Data, scripts, and codes availability

RNA-seq libraries sequenced for this study have been deposited at the European Nucleotide Archive (https://www.ebi.ac.uk/ena/) under accession PRJEB50648. Lists of dataset accessions for each analysis (doi:10.17617/3.XWMBKT) and the 18S rRNA phylogeny (doi:10.17617/3.QLWR38) have been deposited at Edmond. Software and computational workflows have been deposited at Zenodo under the following accessions: doi:10.5281/zenodo.6784075; doi:10.5281/zenodo.6647650; doi:10.5281/zenodo.6647652; doi:10.5281/zenodo.6647679.

## Conflict of interest disclosure

The authors declare that they have no conflict of interest relating to the content of this article.

## Funding

This work was supported by the Max Planck Society. The research leading to these results has received funding from the European Union’s Horizon 2020 research and innovation programme under grant agreement no. 730984, Assemble Plus project.

## Appendix

### Quality metrics of single-cell transcriptome assemblies

Taxonomic composition was variable between samples. Samples with the lowest contamination from non-target taxa were those from cultured isolates, while single-cell environmental samples newly sequenced for this study had consistently about 80% of rRNA reads assigned to the expected target taxon (Figure S1). Furthermore, the proportion of the library composed of SSU rRNA reads was also relatively low in the newly sequenced samples (0.03 to 1.7%). The number of contigs per assembly was highly variable (15389 to 162815), but when only contigs with poly-A tails ≥7 bp were counted, samples from this study had more poly-A tailed contigs (4135 to 13427), compared to assemblies from previously published libraries (48 to 4932) (Figure S2). Presence of poly-A tails can be used to exclude bacterial and rRNA sequences. Contigs with both a poly-A tail and a putatively full-length coding sequence were most abundant for four heterotrich libraries that were prepared from bulk cultured cells instead of single cells (Figure S2, doi:10.17617/3.XWMBKT Table S1). As heterotrichs they were also more closely related to the species used for the reference protein set (*Blepharisma stoltei*).

Different filtering criteria were used to shortlist transcriptome assemblies for prediction of stop codon reassignment to sense vs. prediction of the actual stop codon usage. All ten newly sequenced karyorelict and heterotrich libraries from this study were shortlisted for both analyses. Of the 33 previously published libraries, 15 were used for the former and 16 for the latter (doi:10.17617/3.XWMBKT Table S1).

### Confirmation of phylogenetic identity with 18S rRNA sequences

During collection, each sample was preliminarily identified to a family or genus by morphology under the dissection microscope. Two libraries (N6, N39) were found to be neither karyorelicts nor heterotrichs during the initial screen with phyloFlash (Figure S1). The morphology-based identification of the remaining sequences was verified by a tree of 18S rRNA sequences from the transcriptome assembly (≥80% full length) alongside reference sequences (Figure S3, Table S1). Trachelocercidae spp. (samples N26, N27, N34, N38) could not be identified more specifically to genus, because the taxonomy of several reference sequences were only to family level, and some genera also do not appear monophyletic with 18S rRNA phylogeny [1]. *Remanella* may also be paraphyletic [2] but we chose to retain the name *Remanella* for our samples (N4, N5, N44) because there are only two genera in the family Loxodidae, and the marine species have conventionally been designated *Remanella*.

---

**Figure S1.**
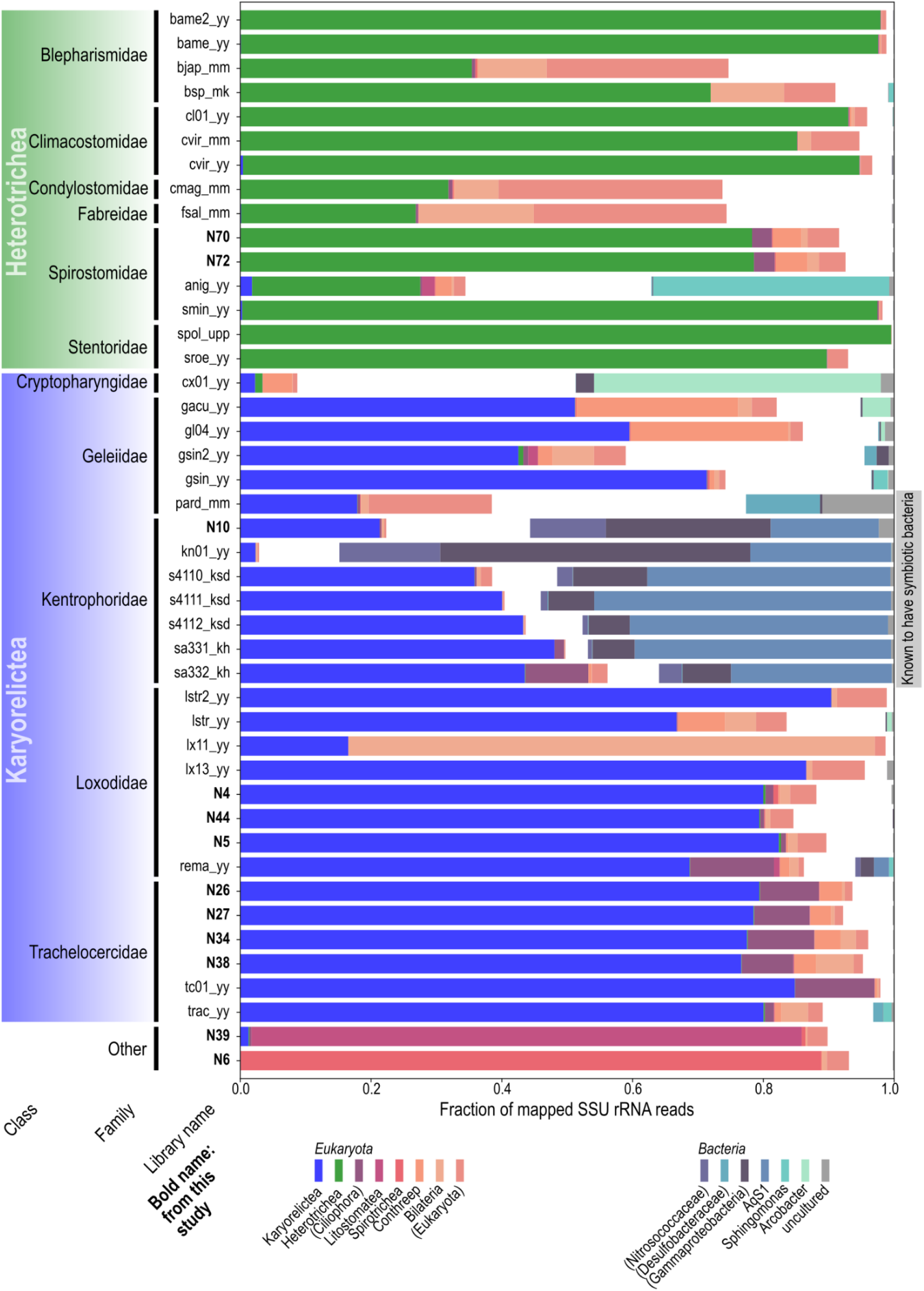
Taxonomic composition of RNAseq libraries, derived from mapping of reads to the SILVA SSU rRNA database, summarized at class level. Only taxa comprising ≥10% of the total in at least one library are shown. Bars representing eukaryotic taxa are aligned to the left, while prokaryotic taxa are aligned to the right.

**Figure S2.**
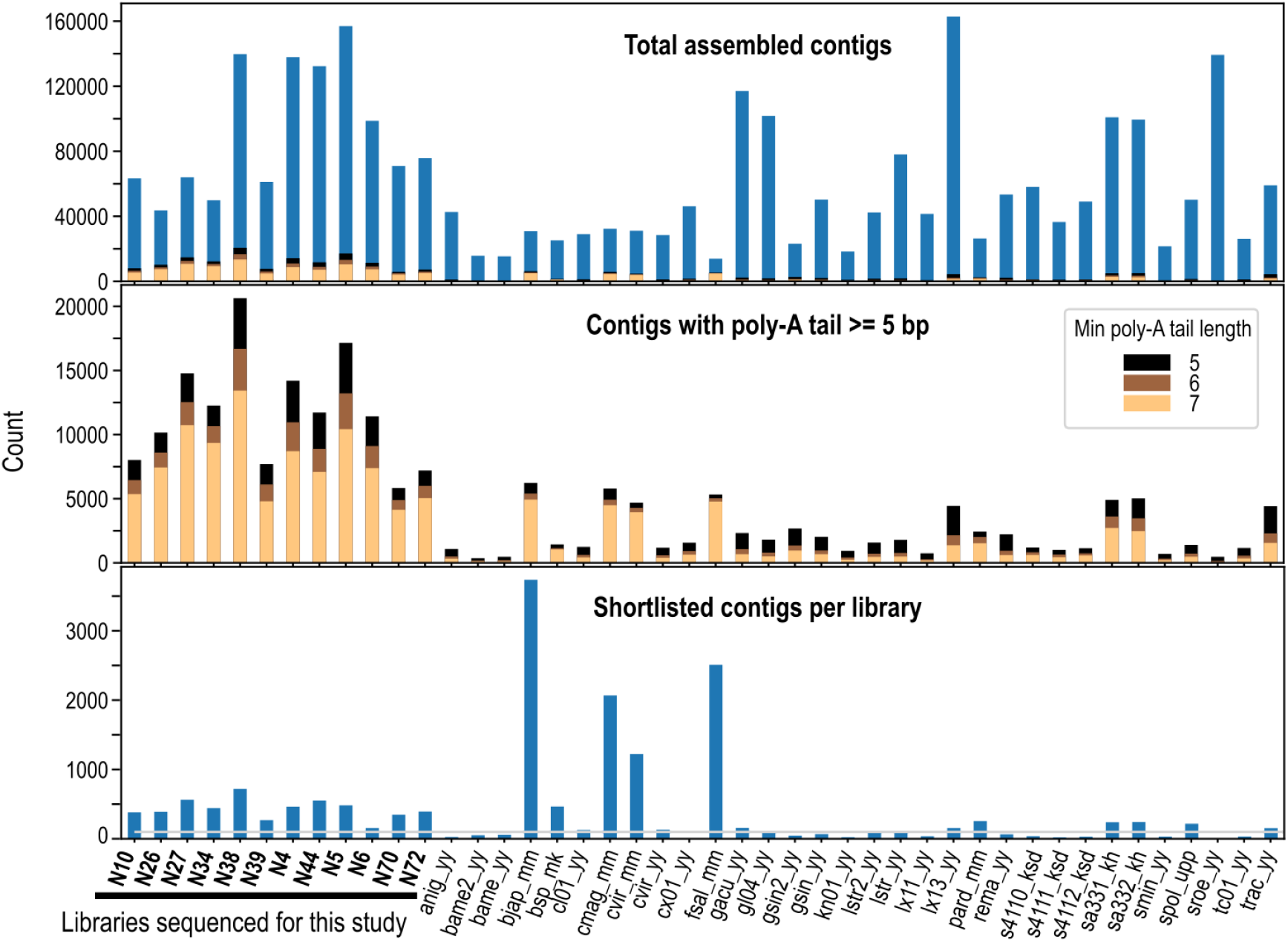
Number of contigs per transcriptome assembly:(top) total number of contigs, (middle) contigs with poly-A-tails, when different minimum lengths were applied, (bottom) putative full-length transcripts with both a poly-A tail (≥7 bp) and >80% Blastx hit vs. reference B. stoltei protein sequence (grey horizontal line: 100 sequences cutoff value).

**Figure S3.**
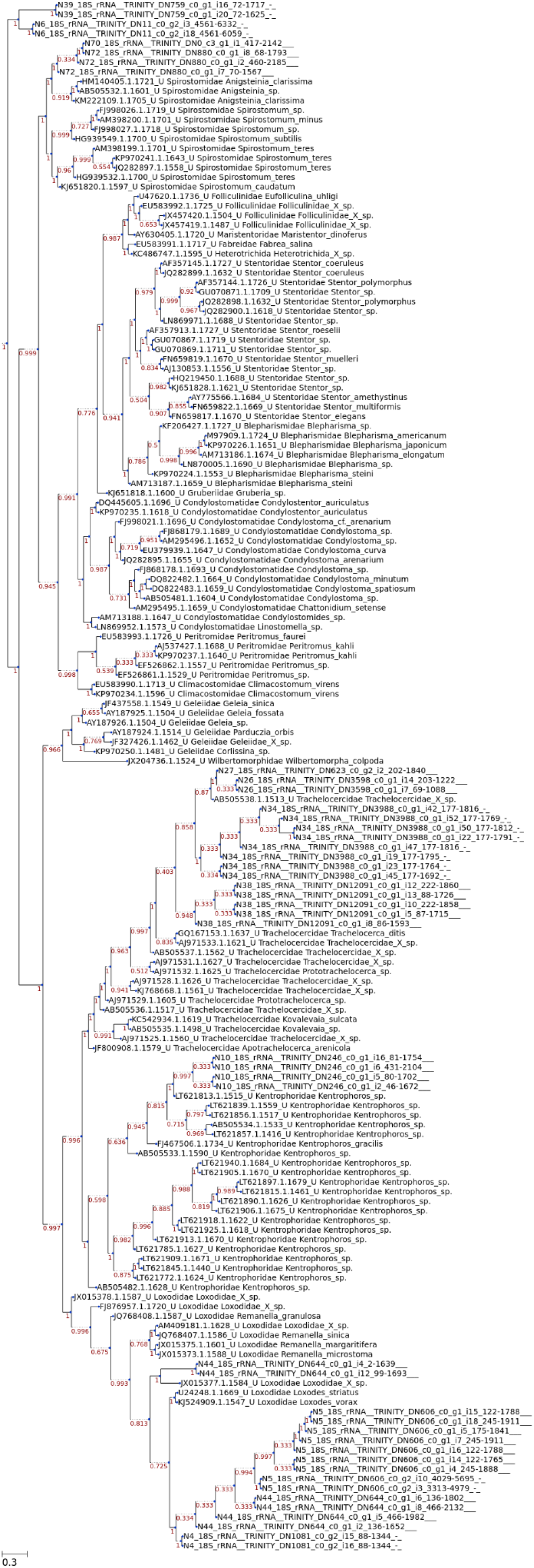
Phylogenetic tree of 18s rRNA sequences from newly sequenced libraries (sequence names beginning with “N”) vs. reference sequences of Karyorelictea and Heterotrichea from the PR2 database (identifier, family, species). Node labels: aBayes support values. Scale bar: Substitutions per site. Dotted lines: Branches spaced to accommodate node labels.

## Notes

https://www.ebi.ac.uk/ena/browser/view/PRJEB50648

https://doi.org/10.17617/3.XWMBKT

https://doi.org/10.17617/3.QLWR38

https://doi.org/10.5281/zenodo.6784075

https://doi.org/10.5281/zenodo.6647650

https://doi.org/10.5281/zenodo.6647652

https://doi.org/10.5281/zenodo.6647679

### Competing Interest Statement

The authors have declared no competing interest.

### Summary of Updates

Added link to peer reviews and recommendation from Peer Community in Genomics; merged supplementary text as appendix to main file; formatting changes

